# Cervical Squamous Intraepithelial Lesions are Associated with Changes in the Vaginal Microbiota of Mexican Women

**DOI:** 10.1101/2021.02.23.432613

**Authors:** ME Nieves-Ramírez, O Partida-Rodríguez, P Moran, A Serrano-Vázquez, H Pérez-Juárez, ME Pérez-Rodríguez, MC Arrieta, C Ximénez-García, BB Finlay

## Abstract

Cervical cancer is an important health concern worldwide and is one of the leading causes of deaths in Mexican women. Previous studies have shown changes in the female genital tract microbe community related to Human Papillomavirus (HPV) infection and cervical cancer, yet this link remains unexplored in many human populations. This study evaluated the vaginal bacterial community among Mexican women with pre-cancerous Squamous Intraepithelial Lesions (SIL). We sequenced the V3 region of the 16S rRNA gene (Illumina Miseq) in cervical samples from 300 Mexican women, including 157 patients with SIL, most of which were HPV positive, and 143 healthy women without HPV infection or SIL. Beta-diversity analysis showed that 14.6% of the variance in vaginal bacterial community structure is related to the presence of SIL. Presence of SIL was also associated with a higher species richness (Chao 1). MaAsLiN analysis yielded independent associations between SIL/HPV status and an increase in the relative abundance *Brachybacterium conglomeratum*, as well as a decrease in *Sphingobium yanoikuyae* and *Lactobacillus* spp. We also identified independent associations between HPV-16, the most common HPV subtype linked to SIL, and *Brachybacterium conglomeratum*. Our work indicates that the presence of SIL and HPV infection is associated with important changes in the vaginal microbiome, some of which may be specific to this human population.

**IMPORTANCE:** HPV plays a critical role in cervical carcinogenesis but is not sufficient for cervical cancer development, indicating involvement of other factors. Vaginal microbiota is an important factor in controlling infections caused by HPV and depending on its composition it can modulate the microenvironment in vaginal mucosa against viral infection. Ethnic and sociodemographic factors influence differences in vaginal microbiome composition, which underlies the dysbiotic patterns linked to HPV infection and cervical cancer across different women populations. Here, we provide evidence for associations between vaginal microbiota patterns and HPV infection, linked to ethnic and sociodemographic factor. To our knowledge, this is the first report of *Brevibacterium aureum* and *Brachybacterium conglomeratum* species linked to HPV infection or SIL.

## INTRODUCTION

Cervical cancer is one of the most common cancers and one of the leading causes of deaths in women worldwide (1, 2). Cervical cancer is causally related with Human Papillomavirus (HPV) infection, an oncogenic virus actively involved in cervical epithelium transformation (3, 4). After HPV infection and persistency, squamous intraepithelial lesions (SIL) development may occur, which may heal or persist and evolve to cancer (1). Despite overwhelming evidence that certain subtypes of HPV are the main causative agents of SIL development and progression to cervical cancer, it is also well-established that HPV alone is not sufficient to induce cervical malignant transformation (4–7). Many factors have been associated with the appearance SIL such as, intermenstrual bleeding, multiparity, use of contraceptives, multiple sexual partners, and smoking (8).

In addition to these variables, it has been proposed that the vaginal microbiota plays an important role in the development of HPV infection leading to cervical neoplasm (9). This is aligned with the endorsed concept in infection biology, in which successful pathogen colonization and infection embodies dynamic interactions between the infecting microbes, host factors and the microbiome (10). The vaginal microbiota is a complex microbial ecosystem influenced by environmental and host factors, as well as ethnic background (11). The vaginal microbiota in healthy women consists of over 200 bacterial species, but this ecosystem is generally dominated by *Lactobacillus* spp. Lactobacilli provide broad spectrum protection by producing lactic acid, bacteriocins and biosurfactants, and by adhering to the mucosa that forms barriers against pathogenic infections in the vaginal microenvironment (2, 12). Upon imbalance of this defense system, physicochemical changes arise, inducing histological alterations of the vaginal mucosa and the cervical epithelium, all of which put a selective pressure on the microbiota (13–15).

Some vaginal microorganisms, such as *Gardnerella, Fusobacteria, Bacillus cohnii, Dialister, Prevotella* and *Mycoplasma*, as well as a decrease in the proportion of *Lactobacillus* spp., have been linked to dysbiosis that would generate an unstable microenvironment, which in turn could enable the effect of key risk factors in cervical cancer (16–19). Some of these changes are responsible for increasing the levels of mucin-degrading enzymes, which may play a role in the degradation of the mucous layer that covers the vaginal and cervical epithelium and endocervical mucus (20, 21). There is evidence of HPV evasion or infection mechanisms that support that microorganisms such as *Sneathia, Anaerococcus, Fusobacterium* and *Gardnerella* are implicated with higher frequency and severity of disease, potentially resulting in pre-cancerous and cancerous cervical lesions (22)

However, these findings are not uniform across studied populations, because, despite the fact that Latin American countries have a high prevalence of HPV and cervical cancer are one of the main causes of death in women in these areas (3, 23–25), including Mexico (7, 9), most of the studies have been conducted in developed countries (26). Likewise, the projected demographic changes in Latin America imply that the current burden of new cervical cancer cases will increase in the next 20 years (2, 27). The evidence observed so far suggests that the ethnic and sociodemographic factors that influence difference in vaginal microbiome composition may also underlie dysbiotic patterns linked to HPV infection and cervical cancer across different Latin America women populations (3,7,9, 23–25). Therefore, there is a growing need for more evidence in Latin America to demonstrate the association between vaginal microbiota patterns and HPV infection and its relationship with the progression of SIL to cervical cancer.

Very little is known about vaginal microbiome differences linked to HPV infection and cervical cancer risk in Latin American women. In this work, we compared the vaginal microbiota in 300 Mexican women with precancerous SIL to healthy controls, while taking into consideration the confounding effect of clinical, behavioral and HPV infection-related variables, and its association with the above-mentioned categorical variables and the condition of HVP infection considering the type of premalignant lesion of cervix and the genetic variants of the virus.

## MATERIAL AND METHODS

### Study design

Healthy women and women infected with HPV regardless of the degree of cervical squamous lesion, over 25 years of age, attending the Instituto Mexicano del Seguro Social (IMSS) in Mexico City were invited to participate as volunteers in this study. Written informed consent was obtained from all volunteers after providing them with detailed information about the study and its characteristics. The clinical research protocol and letter of informed consent were evaluated and approved by the Comité Local de Investigación y de Bioética de la División de Educación e Investigación Médica de la Unidad de Alta Especialidad Médica Pediatría del Instituto Mexicano del Seguro Social (IMSS). All participants completed a study questionnaire that was used to obtain the sociodemographic and risk factor information. Data were registered in a secured database for subsequent statistical analysis.

A total of 300 Mexican women over 25 years old who attended the IMSS from December 2003 to July 2006 were included in this study. These women were divided in two groups: a healthy control group of 143 women with a mean age of 42 years (± 0.65) with three previous Papanicolaou (Pap) tests negative for HPV infection for three consecutive years (a fourth negative Pap result occurred at the time participants were invited to join the study), and diagnosed without SIL, with normal cytology and colposcopy results by the treating gynecologist. The second group (cases) consist of 157 patients with a mean age of 36 years (± 0.89) with different degrees of SIL and result positive for HPV infection based on cytology, histology, and colposcopy examination. This group included women diagnosed with cervical intraepithelial neoplasia from 1 to 3 (CIN1, CIN2 and CIN3) according to the Bethesda classification (28). Participants who had received treatment for vaginal or urinary infections currently, who were pregnant or up to 2 months postpartum, with a history of hysterectomy, or with a severe chronic disease were excluded from the study.

### Samples of vaginal exudate

Samples of vaginal exudate were taken by swabbing the mucosa using sterile Teflon swabs that were placed in a sterile 15 ml conical plastic tube with sterile 0.9 % sodium chloride (Baxter physiological saline solution), the sample was kept at -20 °C until its use for microbiome sequencing analysis.

### Cervical DNA extraction and HPV detection and typing

Cervical DNA was extracted directly from a cervical brushing of each patient. The sample was placed in 1 ml of saline solution at 4 ° C for transport and immediately processed for DNA extraction. DNA was obtained using the proteinase K-SDS lysis technique (29) and was frozen at-20 ° C until use. HPV was detected via PCR, using two sets of oligonucleotides MY09 / MY11 (30) and GP5 / GP6 (31). Cycling conditions were used as previously described for the detection of HPV DNA in cervical cells (30–32). HPV DNA obtained from HeLa cell cultures containing 10 to 50 copies of the HPV-18 ORF L1 was used as a positive control (33). All positive samples for HPV were subsequently typed with the HPVFast 2.0 kit (Pharma Gen SA, Madrid, Spain) according to the manufacturer’s instructions.

### 16S mRNA gene Sequencing

From vaginal DNA samples, the 16S rRNA gene was amplified by PCR in triplicate using bar-coded primer pairs flanking the V3 region as previously described (34). Each 50 ml of PCR mixture contained 22 ml of water, 25 mil of TopTaq master mix, 0.5 ml of each forward and reverse bar-coded primer, and 2 ml of template DNA. The PCR program consisted of an initial DNA denaturation step at 95°C (5 min), 25 cycles of DNA denaturation at 95°C (1 min), an annealing step at 50°C (1 min), an elongation step at 72°C (1 min), and a final elongation step at 72°C (7min). Controls without template DNA were included to ensure that no contamination occurred. Amplicons were run on a 2% agarose gel to ensure adequate amplification. Amplicons displaying bands at ∼160 pb were purified using the illustra GFX PCR DNA purification kit. Purified samples were diluted 1:50 and quantified using PicoGreen (Invitrogen) in the Tecan M200 plate reader (excitation at 480 nm and emission at 520 nm).

For 16S rRNA gene sequencing, each PCR pool was analyzed on the Agilent Bioanalyzer using the high-sensitivity double-stranded DNA (dsDNA) assay to determine approximate library fragment size and verify library integrity. Pooled-library concentrations were determined using the TruSeq DNA sample preparation kit, version 2 (Illumina). Library pools were diluted to 4 nM and denatured into single strands using fresh 0.2 N NaOH. The final library loading concentration was 8 pM, with an additional PhiX spike-in of 20 %. Sequencing was carried out using a Hi-Seq 2000 bidirectional Illumina sequencing and cluster kit, version 4 (Macrogen, Inc.). PCR products were visualized on E-gels, quantified using Invitrogen Qubit with PicoGreen, and pooled at equal concentrations, according to a previous report (35).

### Bioinformatic analysis of 16S rRNA gene sequences

All sequences were processed using Mothur according to the standard operating procedure as previously described (36). Quality sequences were obtained by removing sequences with ambiguous bases, a low-quality read length and/or chimeras identified using chimera uchime. Quality sequences were aligned and compared to the SILVA bacterial references alignment and OTUs were generated using a dissimilarity cutoff of 0.03. The sequences were classified using the classify seqs command.

### Statistical Analysis

Differences in frequencies for categorical variables between cases and controls were evaluated using the chi squared. Risk was estimated and expressed as an odds ratio (OR) and a 95% confidence interval (CI). For numerical variables the Mann-Whitney or Student t tests were used based on the D’Agostino & Pearson normality test. We assessed the vaginal microbial diversity and the relative abundance of bacterial taxa using Phyloseq (37) along with additional R-based computational tools in R-studio (R-Studio, Boston, MA). Principal components analysis (PCA) was conducted using Phyloseq and statistically confirmed by PERMANOVA (Adonis test). The Chao 1 and Shannon diversity indices were calculated using Phyloseq and statistically confirmed by Mann-Whitney (GraphPad Prism software, version 5c, San Diego, CA). Lefse analysis (38, 39) was used to evaluate OTU-level microbiome differences between cases and controls. Multivariate association with linear models (MaAsLin, (38)) were used to calculate differentially abundant OTUs between the cases and controls, including several other study variables available from the metadata. The following covariates were fitted into the MaAsLin model based on previously reported associations with HPV infection or with microbiome shifts: SIL grade, HPV infection, HPV type, smoking, intermenstrual bleeding, sexual activity status, use of contraceptives, type of contraceptive, genital hygiene, age, age of sexual debut, number of sexual partners, number of sexual partners by age, number of pregnancies, number of births and number of miscarriages. The random forest classifier in R was applied to determine if differential microbiome taxa would be discriminant between cases and controls.

## RESULTS

### Study participants characteristics: Cases vs. controls

A total of 300 samples were analyzed, 143 controls 157 cases. Of the 157 cases, 112 were diagnosed with low squamous intraepithelial lesion (LSIL) (women diagnosed with HPV infection and cervical intraepithelial neoplasia 1 (CIN 1), and 45 were diagnosed with HPV infection and high squamous intraepithelial lesion (HSIL) (women diagnosed with CIN 2 or CIN 3). All women were cancer free. For the selection of participants, HPV infection was determined by positive cytological, histological and colposcopy analysis.

However, by molecular analysis, within cases, the frequency of positivity to HPV infection detected was 90.45%, of which HPV-16, -58 and -18 types were the most frequently detected with 49.04%, 14.65% and 10.83%, respectively. Most of the women in both groups did not smoke (75.16% -cases vs 70.63% controls), had a regular menstrual period (70.70% cases vs 69.23% controls), and most do not have intermenstrual bleeding (82.80% cases vs 89.51% controls). Statistically significant differences between groups were detected in relation with active sexual life at the time of the study (75.16% cases vs 92.31% controls), use of contraceptives (66.24% cases vs 53.15% controls) in the control group (P = 0.021), and genital hygiene, recorded by the frequency of vaginal douching (83.44% cases group vs 53.85% control group) and such differences were statistically significant (P < 0.0001). More details of the characteristics of each group are described in Table 1. When comparing continuous variables, cases and controls differed by age (36.3±0.9 cases vs 42.9±0.7 controls), number of sexual partners by age (0.0038 cases vs 0.028 controls) and number of miscarriages (0.01 cases vs. 0.014 controls; Table 2).

**Table 1.** Characteristics of Study Population. Categorical Variables.

| Variable | Subcategory | With SILs<br>(N=157) | Without SILs<br>(N=143) | P | OR | CI |
| --- | --- | --- | --- | --- | --- | --- |
| SILs Grade | Low grade | 112 | 0 | N/A | N/A | N/A |
|  | High grade | 45 | 0 |  |  |  |
| HPV | Positive | 142 (90.45%) | 0 | <0.0001 | ∞ | 355.4<br>to ∞ |
|  | Negative | 15 (9.55%) | 143 (100%) |  |  |  |
| HPV type | HPV-16 | 77(49.04%) | 0 | N/A | N/A | N/A |
|  | HPV-58 | 23(14.65%) | 0 |  |  |  |
|  | HPV-18 | 17(10.83%) | 0 |  |  |  |
|  | HPV-31 | 7(4.46%) | 0 |  |  |  |
|  | HPV-11 | 4(2.55%) | 0 |  |  |  |
|  | Other | 14(8.92%) | 0 |  |  |  |
|  | HPV Neg | 15(9.55%) | 143(100%) |  |  |  |
| Smoking | Yes | 39(24.84%) | 42(29.37%) | 0.38 | 0.79 | 0.47<br>to<br>1.32 |
|  | No | 118(75.16%) | 101(70.63%) |  |  |  |
| Menstrual period | Regular | 111(70.70%) | 99(69.23%) | 0.78 | 1.07 | 0.66<br>to<br>1.73 |
|  | Irregular | 46(29.30%) | 44(30.77%) |  |  |  |
| Intermenstrual<br>bleeding | Yes | 27(17.20%) | 15(10.49%) | 0.095 | 1.77 | 0.89<br>to<br>3.41 |
|  | No | 130(82.80%) | 128(89.51%) |  |  |  |
| Sexually active (at<br>study assessment) | Yes | 118(75.16%) | 132(92.31%) | <0.0001 | 0.25 | 0.12<br>to<br>0.51 |
|  | No | 39(24.84%) | 11(7.69%) |  |  |  |
| Use of<br>contraceptive(s) | Yes | 104(66.24%) | 76(53.15%) | 0.021 | 0.58 | 0.37<br>to<br>0.93 |
|  | No | 53(33.76%) | 67(46.85%) |  |  |  |
| Contraceptive type | IUD | 39(24.84%) | 17(11.89%) | 0.0001 | N/A | N/A |
|  | Tubal ligation | 27(17.20%) | 21(14.69%) |  |  |  |
|  | Hormonal | 20(12.74%) | 13(9.09%) |  |  |  |
|  | Condom | 10(6.37%) | 3(2.10%) |  |  |  |
|  | IUD+Tubal<br>Ligation | 2(1.27%) | 0 |  |  |  |
|  | Other | 6(3.82%) | 10(6.99%) |  |  |  |
|  | Did not specify | 0 | 13(9.09%) |  |  |  |
|  | None | 53(33.76) | 66(46.15%) |  |  |  |
| **Vaginal douching** | Yes | 26(16.56%) | 66(46.15%) | **<0.0001** | 0.23 | 0.14<br>to<br>0.39 |
|  | No | 131(83.44%) | 77(53.85%) |  |  |  |

P values in bold denote statistical significance (P>0.05)

**Table 2.**
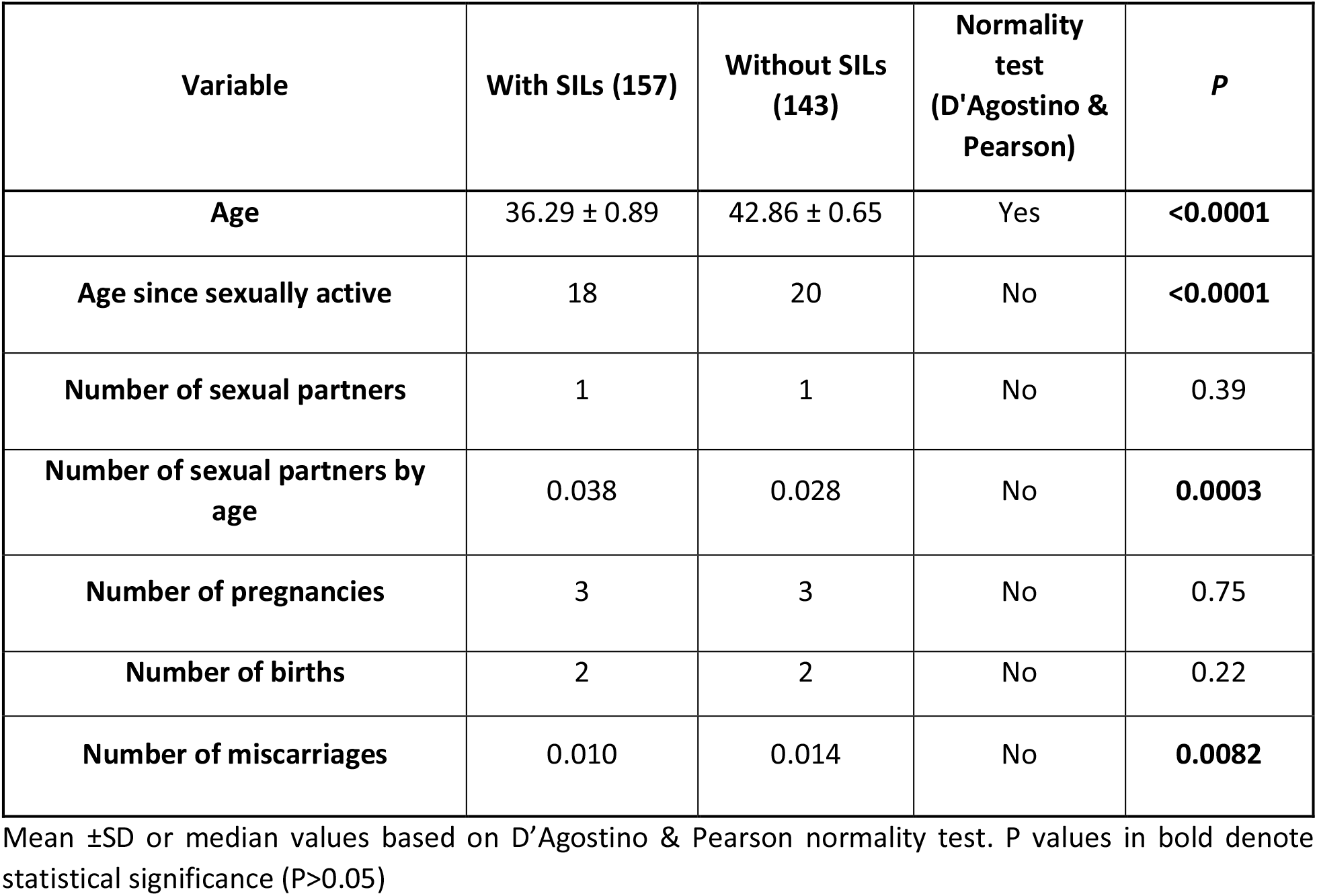
Characteristics of Study Population. Numerical variables.

### Associations between the Vaginal Microbiota SIL status

We determined the bacterial community by amplification and sequencing of the 16S rRNA gene (V3 region). The presence of SIL was associated with changes in bacterial alpha and beta diversity (Figure 1), with notable compositional differences at the family and genus level (Figure 2). Beta-diversity analysis, measured by Principal Component Analysis (PCoA; Bray Curtis distance, Figure 1A) indicated that cervical SIL explain 14.6% of the variation in vaginal bacterial community structure (N=300; Adonis P>0.001). Presence of SIL was also associated with significantly higher species richness than women without SIL (Chao1; P=2.78e-07; Fig.1B). Only a trend for an increase in alpha diversity (Shannon index) was observed in SIL-positive participants, suggesting that the broadest diversity change is explained by bacterial community richness.

**FIGURE 1.**
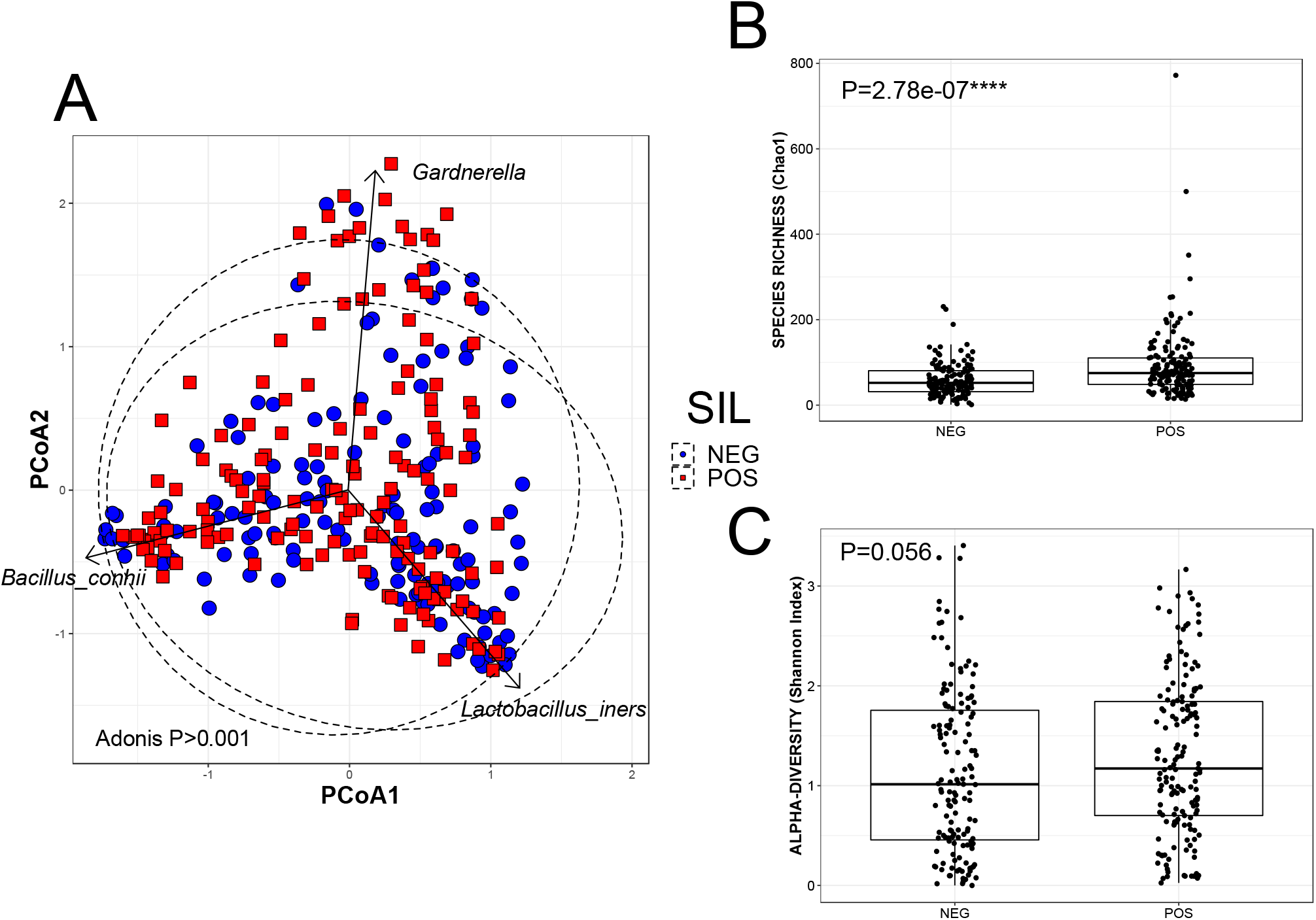
(A)Principal component analysis (PCoA) ordination of variation in beta-diversity of human cervical bacterial communities in adult Mexican women based on Bray-Curtis dissimilarities. Color and shape represent presence of squamous cervical intraepithelial lesions (SIL); blue circles represent absence of SIL and red squares represent presence of SIL. PERMANOVAs indicate the SIL represent 14.6% of the variation in vaginal bacterial community structure (N=300; Adonis P>0.001). Arrows represent loading plot coordinates for the three most abundant OTU features in the dataset. Variation in **(B)** species richness (Chao1) and **(C)** alpha-diversity (Shannon index) of vaginal bacterial communities between women with (POS) and without (NEG) SIL. Stars denote statistical significance (N=300; Kruskal-Wallis test; Chao1 P=2.78e-07).

**FIGURE 2.**
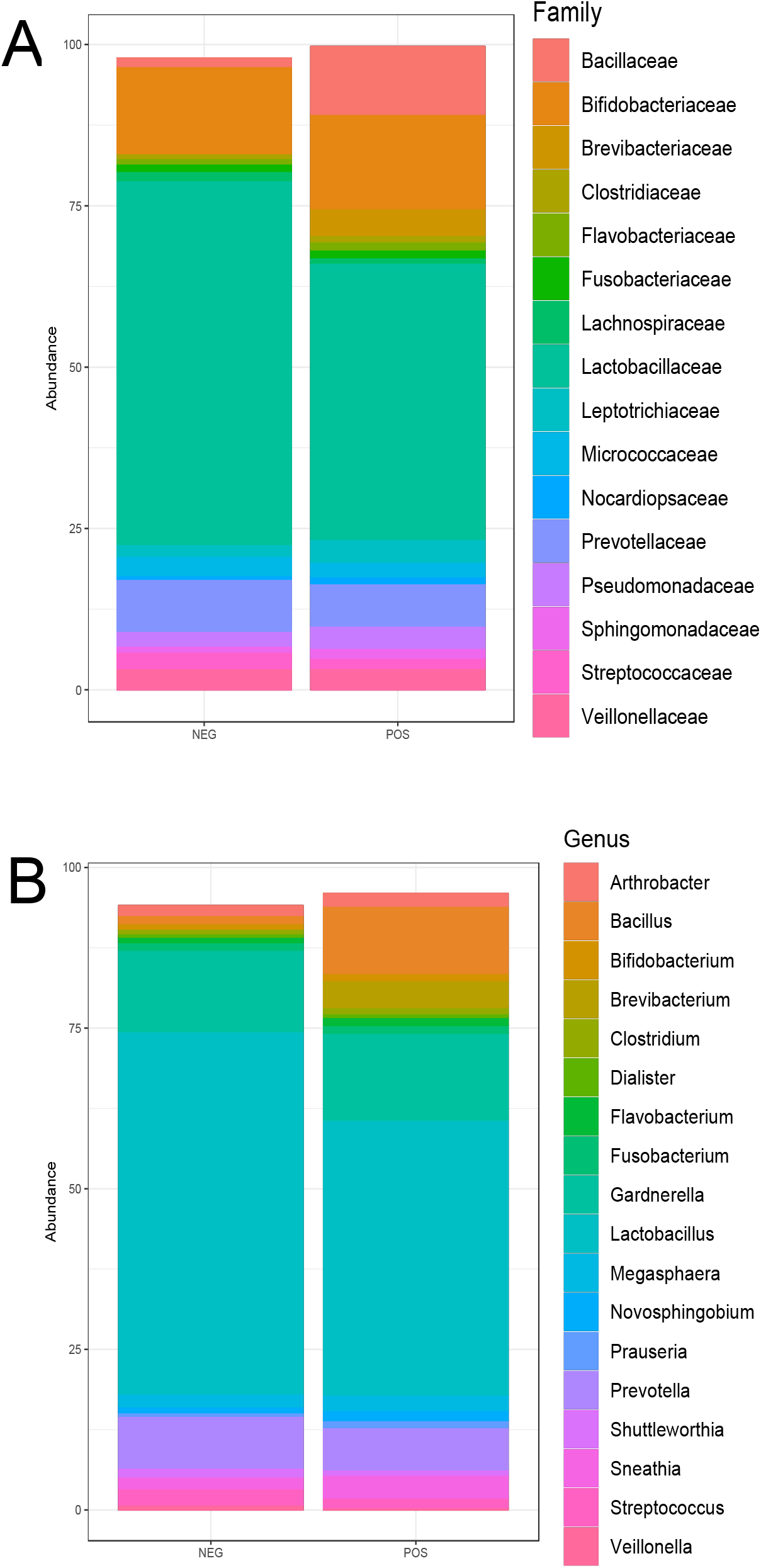
Variation in taxonomic composition of vaginal bacterial communities at the family (A) and genus (B) levels between women with and without cervical SIL (N=300).

We utilized Lefse to identify OTU-level difference between SIL positive and negative groups. In this model, features are first tested to determine if they are differentially distributed. Microbial features violating the null hypothesis are further analyzed in a secondary analysis, in which a Latent Dirichlet Allocation (LDA) model is built to detect and rank microbiome feature differences among groups. Lefse identified a greater abundance of 12 OTUs in SIL-positive women, with OTU 14 (S_*Brevibacterium*_*aureum*), OTU 117 (F_Veillonellaceae), OTU 28 (S_*Brachybacterium*_*conglomeratum*), and OTU 101 (*Lactobacillus iners*) as the most differentially abundant features (Figure 3). In contrast, OTU 62 (*Sphingobium yanoikuyae*), OTU 129 (*Zoogloea* sp.) and OTU 80 (*Sphingobium* sp.) were detected in higher abundance in the control samples (Figure 3). Among these, *Brevibacterium aureum* was exclusively detected in cases (Figure 4A), whereas *Zoogloea* sp. was exclusively detected in controls (Figure 4B). Other taxa that reached almost exclusive detection in either group include *Brachybacterium conglomeratum* and *Prevotella* sp. (Figure 4). Given this, we evaluated if any of these features could be used to predict SIL status by applying Random Forest analysis, which showed that none of the features can accurately classify a participant in the SIL positive or negative groups (overall error rate=0.67).

**FIGURE 3.**
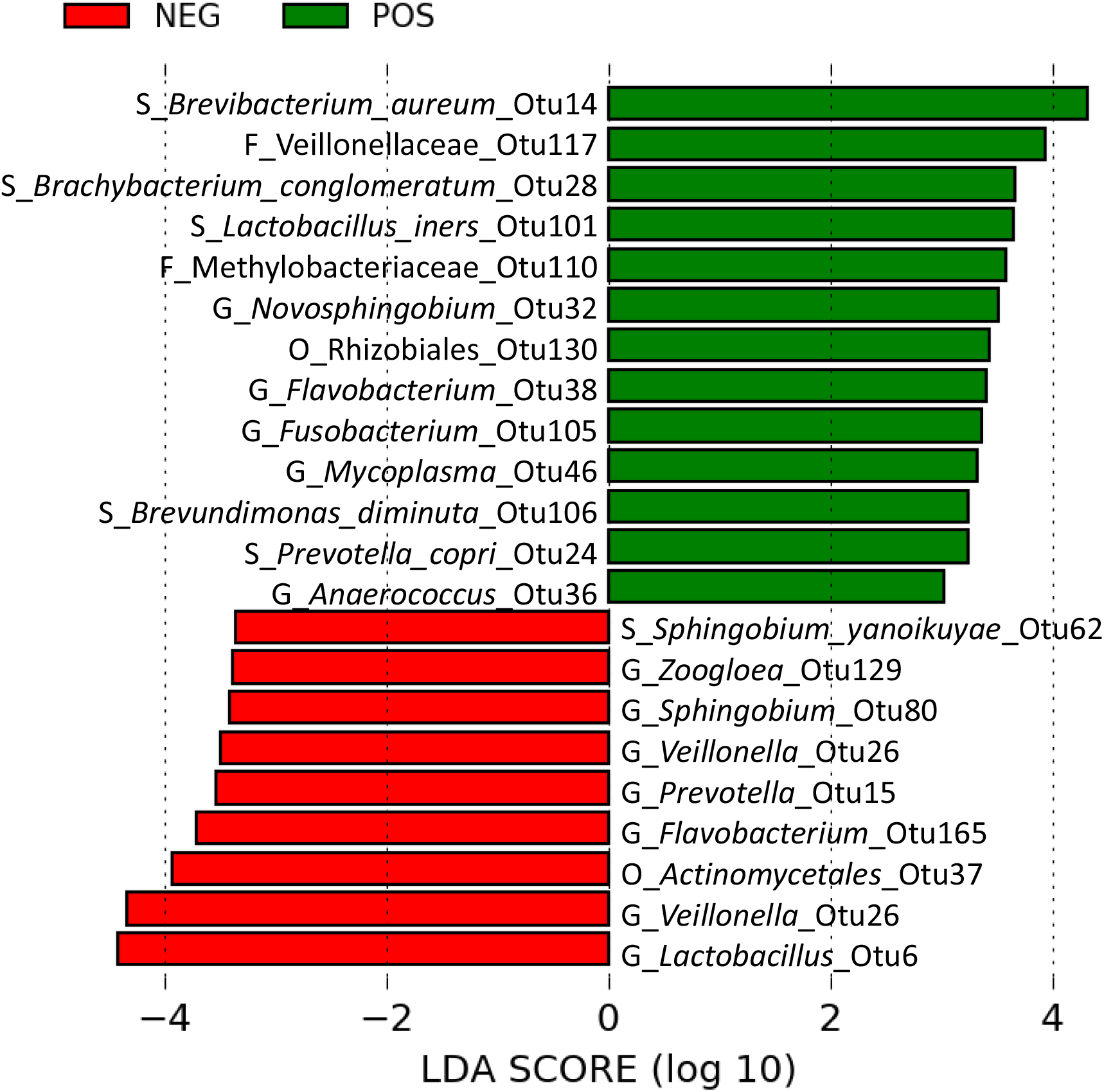
Differentially abundant taxa (OTU-level) in women with (green) or without (red) cervical SIL, identified by linear discriminant analysis (LDA). Only taxa meeting an LDA significant threshold >2 are shown (N=300; Lefse (39)).

**FIGURE 4.**
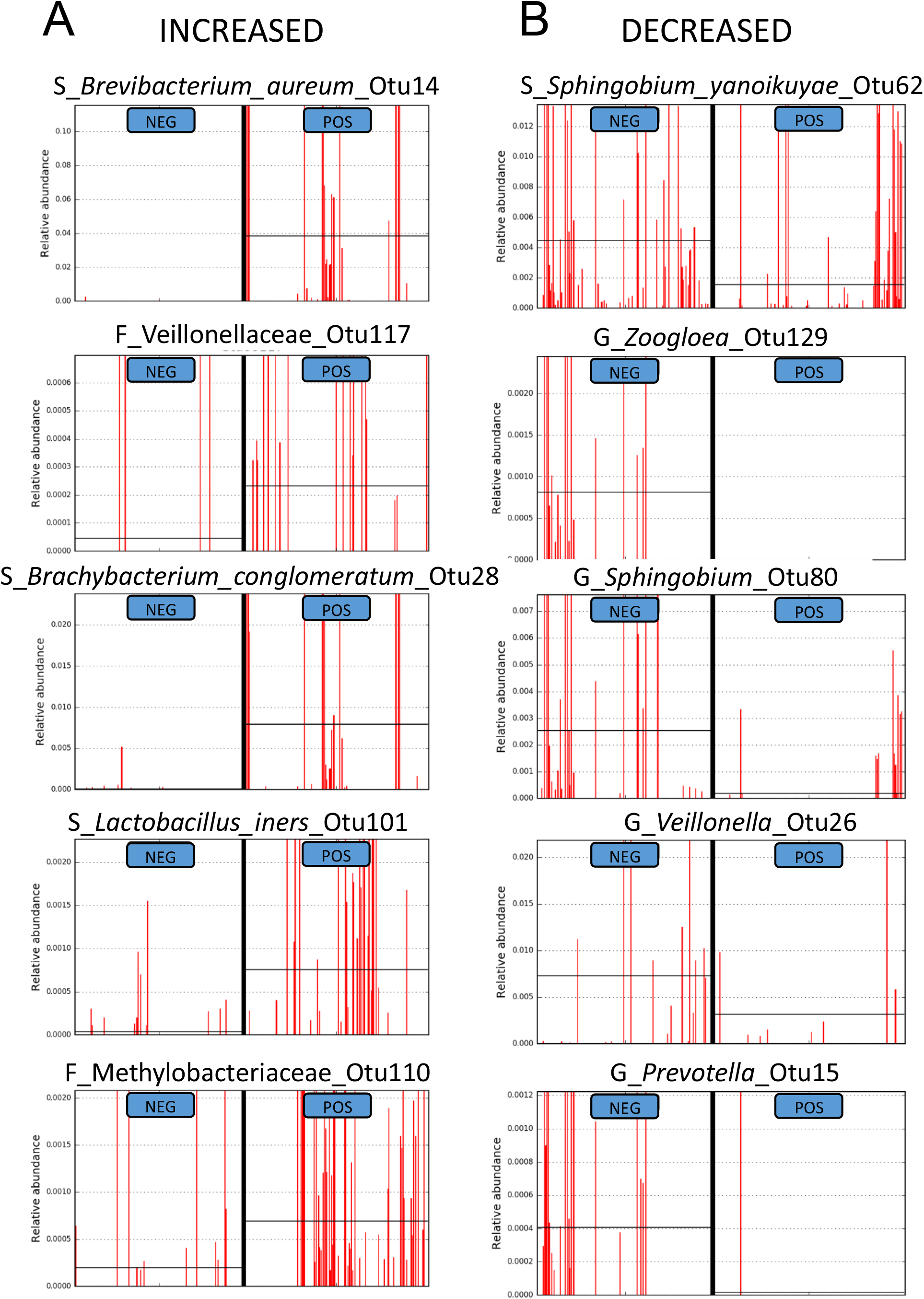
Histogram of most discriminant increased (A) or decreased (B) OTUs in women with (POS) and without (NEG) cervical SIL. Five features were chosen per category, based on effect size calculated by LDA (N=300; Lefse (39)). Red lines indicate relative abundance for each sample, and horizontal black line denotes median value.

Given the importance to control for potential confounding variables, including several collected in this study that could explain or correlate with the detected associations between SIL status and the microbiota, we utilized MaAsLin. MaAsLin is a multivariate linear modeling tool with boosting that tests for associations between specific microbial taxa and continuous and/or Boolean metadata. This method reduces the total amount of correlations to be tested, therefore allowing for improvements in the robustness of the additive general linear models. With MaAsLin, we found significant independent associations between SIL positive status and *Branchybacterium conglomeratum*, as well as between SIL negative status and *Lactobacillus* sp. and *Sphingobium yanoikuyae*. This indicates that no other variable explained the taxonomic differences observed SIL status and these bacterial taxa. Interestingly, other independent associations were also detected between HPV subtypes or contraception use and several bacterial taxa (Table 3).

**Table 3.** Differential OTUs in relation to study variables (MaAsLiN). Features organized in ascending order of adjusted P values

| Variable | Feature | Value | P.value | Q.value |
| --- | --- | --- | --- | --- |
| Contraception | *G_Fusobacterium*_Otu105 | IUD | 2.4E-110 | 7.8E-106 |
| HPV_type | *G_Mycoplasma*_Otu46 | HPV-90 | 3.2E-39 | 4.2E-35 |
| Contraception | *S_Acinetobacter_lwoffii*_Otu127 | IUD | 7.3E-31 | 8.1E-27 |
| Contraception | *F_Tissierellaceae*_Otu174 | IUD | 3.8E-15 | 3.2E-11 |
| Contraception | *S_Brevundimonas_diminuta*_Otu106 | IUD | 2.5E-13 | 1.7E-09 |
| Contraception | *F_Micrococcaceae*_Otu49 | IUD | 3.9E-12 | 2.2E-08 |
| SILs | *F_Brachybacterium_conglomeratum*_Otu28 | POS | 6.5E-11 | 3.1E-07 |
| SILs | *G_Lactobacillus*_Otu6 | NEG | 1.2E-08 | 3.3E-05 |
| SILs | *S_Sphingobium_yanoikuyae*_Otu62 | NEG | 2.0E-07 | 3.1E-03 |
| Contraception | O_BD7-3_Otu216 | IUD | 5.70E-07 | 1.6E-04 |
| SILs | *G_Lactobacillus*_Otu23 | NEG | 2.42E-06 | 4.14E-04 |
| HPV_type | *S_Brachybacterium_conglomeratum*_Otu28 | HPV-16 | 1.02E-05 | 1.93E-02 |
| HPV_type | *S_Lactobacillus_iners*_Otu101 | HPV-83 | 1.69E-05 | 2.8E-02 |
| Age | *S_Streptococcus_anginosus*_Otu33 | Age | 1.71E-04 | 2.5E-02 |

## DISCUSSION

Several factors are known to play a role in cervical carcinogenesis, with HPV infection being one of the most important in the development of the disease (1). There are more than 100 types of HPV, of which at least 14 high-risk HPV types have been defined as carcinogenic (40). In this study we found that more than 90% of the group of cases were HPV positive and that almost 50% of HPV infections are caused by the HPV-16 type, followed by HPV-58 and -18, all of them considered as high-risk HPVs worldwide (41). This predominance of the HPV-16 type was expected since it is generally accepted that HPV-16 is the major high-risk genotype in Mexico and in the world (42, 43). We also found HPV-58 as the second most prevalent genotype, in 14.65% of the cases, aligned with has been reported in Asia (14.36 – 15.90 %) (42, 43).

Our study also revealed several other factors associated with SIL status, some of which reaffirm previously reported links (44). Factors positively associated with SIL included younger age, HPV infection, younger age of sexual debut, number of sexual partners by age, number of pregnancies and births by age, and the use of contraceptives, with the biggest difference explained by IUD use. In contrast, being sexually active at the time of the study, vaginal douching and number of miscarriages were linked to a reduced risk to SIL in this group of women.

Regarding contraceptive use, our result differs from that reported by Cortessis *et al*. (45), in which they indicated that invasive cervical cancer can be approximately 30% less frequent in women who have used IUD. Likewise, Agenjo *et al*. (46) described an inverse relationship between IUD use and cervical cancer risk, with women using IUD reporting half the risk of developing this type of cancer. Our contrasting results, however, are in line with previous microbiome correlations with cervical cancer. We found significant correlations with IUD use and the presence of *Acinetobacter lwoffii*, which has been previously reported in HPV-positive women (47). In addition, we detected an independent positive correlation with the use of IUD and *Fusobacterium* sp. and a taxon of the Tissierellaceae family. *Fusobacterium* has been studied as a possible diagnostic biomarker of cervical cancer since it is positively correlated with tumor differentiation (48). Furthermore, both Tissierellaceae and Fusobacteriaceae have been reported as the most abundant microorganisms in cervical carcinoma (49). Thus, while the relationship between IUD use and cervical cancer remains varied across studies, our results support that IUD use is linked to vaginal bacteria previously detected in greater abundance in cervical cancer. The fact that we detected a link between contraception and SIL for IUD only, and not for other forms of hormonal or physical contraception methods may suggest that the use of IUD could favor the growth of specific bacterial species that may either induce changes in the cervical microenvironment that could favor HPV infection, or alternatively, facilitate HPV infection via microbial interactions. It is also possible that these bacterial changes are a consequence of the anatomical and immune changes associated with SIL and cervical cancer. Future work should study host-microbe interactions involving these bacterial species and HPV in experimental models of cervical cancer, as well as microbiome features associated with IUD use in healthy women. This mode of contraception is widely used by women across the world; thus, it is important to further elucidate if microbial species linked to IUD use could be causally linked to HPV infection and cervical cancer risk.

While it is unclear why younger age was linked to SIL in our study, it is likely that it relates to the common age of onset of SIL, which occurs between 25-35 years of age (50, 51). In contrast, healthy women would be less likely to visit the IMSS for a routine gynecological visit. Our microbiome results did not find any differences associated with age, suggesting that age did not confound our results. Several study variables linked sexual activity with SIL, included younger age of sexual debut and number of sexual partners per age. These and other related sexual behavioral factors have been previously linked with SIL, HPV infection and cervical cancer risk (52). Interestingly, our study revealed that vaginal douching was linked to a reduced risk of SIL (OR 0.23 CI 0.14-0.39). Studies on cervical cancer and vaginal douching have reported positive, negative and no associations (53). Although it is unlikely that SIL would lead to symptoms that would motivate genital douching, this practice is more common among women with other risk factors linked to sexually transmitted infections, which are a common cause of symptoms.

Among the predominant components of a healthy vaginal microbiome are *Lactobacillus* species, including *L. crispatus, L. iners, L. jensenii*, and *L. gasseri* (17, 54), which results in reduced community diversity. Indeed, bacterial richness increases as *Lactobacillu*s spp. levels are reduced in association with precursor lesions of cervical cancer (17) and with HPV infection itself (2, 55, 56). In support to this, our results showed higher species richness in cases as well as shifts on beta-diversity. Compositional differences involved several taxa, including lactobacilli. While one *L. iners* OTU had greater relative abundance in positive cases, two other significantly more predominant *Lactobacillus* OTUs were decreased in women with SIL, explaining on overall reduction in lactobacilli (Figure 2). *L. iners* has been previously associated with a dysbiotic community and displays a series of characteristics that make this species different from other known vaginal lactobacilli (57–59). For instance, *L. iners* is a lower producer of D-lactic acid and induces IL-8 secretion causing pro-inflammatory activity in the cervix, which may influence the progression of cervical intraepithelial neoplasia (15). In other studies, the dominance of *L. iners* and interactions with other vaginal anaerobic microorganisms alters the balance of the vaginal microbiota in association with cervical intraepithelial neoplasia (13).

The most discriminant microbial differences between cases and controls involved *Brevibacterium aureum* and *Brachybacterium conglomeratum* (increased in cases), as well as *Zoogloea* sp. and *Prevotella* sp. (increased in controls; Figure 4). While these differences were very significant, these species were not uniformly present among either group suggesting that interindividual compositional differences may prevent to identify microbiota species with biomarker potential for HPV infection or SIL. However, our study identified *Brachybacterium conglomeratum* as independently associated with SIL and with HPV-16, the most common subtype detected in our study. This prompts for future investigation on the link of this bacterial species with SIL risk associated with this specific HPV subtype and raises the possibility that microbiome links with HPV infection are subtype specific. To our knowledge, this is the first time this species is linked to HPV infection or SIL. *B. conglomeratum* has not been readily reported in vaginal microbiome studies either, which have mainly surveyed North American and European populations (60–62). This finding underlines the importance to consider ethnicity and geography-driven differences in human microbiome studies, as dysbiotic patterns may be population-specific.

## ACKNOWLEDGMENTS

The present work was supported by PAPIIT-DGAPA-UNAM grant IN215018 from the Universidad Nacional Autónoma de México (UNAM) and grant number -272601 from Consejo Nacional de Ciencia y Tecnología (CONACyT), as well as Canadian Institutes of Health Research (CIHR) grants to B.B. We appreciate the technical support of M in Sc. Martha Elena Zaragoza and QFB Angeles Padilla.

